# Construction and analysis of mRNA, miRNA, lncRNA, and TF regulatory networks reveal the key genes in prostate cancer

**DOI:** 10.1101/323543

**Authors:** Su-Liang Li, Yun Ye, Sheng-Yu Wang

**Author notes:** These authors contributed equally to this study (Su-Liang Li, Yun Ye). Address reprint requests to: Yun Ye Department of Clinical Laboratory, The First Affiliated Hospital of Xi’an Medical University, No. 48, West Fenghao Road, Xi’an, Shaanxi Province 710077, People’s epublic of China.

## Abstract

***Purpose*:** Prostate cancer (PCa) causes a common male urinary system malignant tumour, and the molecular mechanisms of PCa remain poorly understood. This study aims to investigate the underlying molecular mechanisms of PCa with bioinformatics.

***Methods***: Original gene expression profiles were obtained from the GSE64318 and GSE46602 datasets in the Gene Expression Omnibus (GEO). We conducted differential screens of the expression of genes (DEGs) between two groups using the R software limma package. The interactions between the differentially expressed miRNAs, mRNAs and lncRNAs were predicted and merged with the target genes. Co-expression of the miRNAs, lncRNAs and mRNAs were selected to construct the mRNA-miRNA and-lncRNA interaction networks. Gene Ontology (GO) and Kyoto Encyclopaedia of Genes and Genomes (KEGG) pathway enrichment analyses were performed for the DEGs. The protein-protein interaction (PPI) networks were constructed, and the transcription factors were annotated. The expression of hub genes in the TCGA datasets was verified to improve the reliability of our analysis.

***Results***: The results demonstrated that 60 miRNAs, 1578 mRNAs and 61 lncRNAs were differentially expressed in PCa. The mRNA-miRNA-lncRNA networks were composed of 5 miRNA nodes, 13 lncRNA nodes, and 45 mRNA nodes. The DEGs were mainly enriched in the nuclei and cytoplasm and were involved in the regulation of transcription, related to sequence-specific DNA binding, and participated in the regulation of the PI3K-Akt signalling pathway. These pathways are related to cancer and focal adhesion signalling pathways. Furthermore, we found that 5 miRNAs, 6 lncRNAs, 6 mRNAs and 2 TFs play important regulatory roles in the interaction network. The expression levels of EGFR, VEGFA, PIK3R1, DLG4, TGFBR1 and KIT were significantly different between PCa and normal prostate tissue.

***Conclusion*:** Based on the current study, large-scale effects of interrelated mRNAs, miRNAs, lncRNAs, and TFs were revealed and a model for predicting the mechanism of PCa was provided. This study provides new insight for the exploration of the molecular mechanisms of PCa and valuable clues for further research.

---

Prostate cancer (PCa) involves a common male urinary system malignant tumour that has the highest incidence among European and American populations [1, 2]. This disease not only seriously affects the quality of life of patients but is also associated with financial burdens for the society and family [3, 4]. PCa is closely regulated by various cytokines, related genes and intercellular signalling networks in the course of the development of prostate cancer. However, the molecular mechanisms remain poorly understood. Therefore, it is important to study the molecular mechanisms of prostate cancer and to prevent the disease and develop effective treatments for the disease.

MiRNAs are a class of non-coding small RNAs that can be combined with the 3-’UTR of target mRNA to regulate gene expression, which leads to abnormal expression of the target genes. A large number of studies have reported abnormal expression of miRNAs in prostate cancer that involve multiple miRNAs, such as miR-16, 21, 221, 375, 34a, 141, and let-7a. The abnormal expression of miRNAs has been confirmed to be closely related to long non-coding RNAs (lncRNAs) and transcription factors (TFs). lncRNAs regulate the expressions of miRNAs by binding and sequestering the target miRNAs and participating in the expression regulation of mRNAs [5, 6]. As an important factor in gene transcription and post-transcription regulation, TFs are also involved in control with miRNAs [7, 8]. However, further research is needed to determine the regulation mechanisms of miRNAs, lncRNAs, TFs and mRNAs for PCa.

Microarray analysis can quickly identify all of the genes that are expressed at the same time-point [9]. Future research could benefit from the integration and analysis of the data [10]. In this work, we identified differentially expressed genes (DEGs) in prostate cancer from the GSE64318 and GSE46602 datasets. We performed Gene Ontology and signalling pathway enrichment analyses for differentially expressed miRNAs targeting mRNAs. Furthermore, we analysed the mRNAs-miRNAs-lncRNAs and protein-protein interactions (PPIs) network to reveal the interactions and identified some factors that may be associated with regulatory mechanisms in PCa. Finally, we analysed the hub genes based on PPI network and TCGA datasets. This study will contribute to the exploration of the molecular mechanism of prostate cancer and provide valuable clues for further research.

## Material and methods

### Raw data

The datasets used in the present study were downloaded from the National Center of Biotechnology Information (NCBI) Gene Expression Omnibus (GEO) (https://www.ncbi.nlm.nih.gov/geo/) [11]. The experimental articles that were used to compare the RNAs in the prostate cancer tissue and noncancerous prostate tissue from patients were included. The original gene expression profiles were obtained from GSE64318 dataset 26089375 and GSE46602 dataset 26522007. The GSE64318 dataset includes microRNA expression profiles of the prostate biopsy samples from 54 prostate biopsy specimens (tumours and normal tissues). The platforms used in these data are the GPL8227 Agilent-019118 Human miRNA Microarray 2.0 G4470B (miRNA ID version). GSE46602 dataset, which includes genome expression profiling of tumour tissue specimens from 36 patients with prostate cancer and normal prostate biopsies from 14 patients. The platform used to analyse these data was the GPL570 Affymetrix Human Genome U133 Plus 2.0 Array.

### Identification of differentially expressed genes

We used the robust multi-array average algorithm to perform background correction and quartile data normalization of the downloaded data [12]. Probes without a corresponding gene symbol were then filtered, and the average value of the gene symbols with multiple probes was then calculated. Student’s *t*-tests and fold-change (FC) filtering were conducted to screen for the differentially expressed genes (DEGs) between the two groups using the R software limma package [13]. With a threshold P-value <0.05 and an absolute value of FC> 2, volcano plot filtering was performed using the R software ggplot2 package to identify the DEGs with statistical significance between the two groups. Hierarchical clustering and combined analyses were performed for the DEGs.

### Prediction of the mRNA-miRNA-lncRNA interactions

The interactions between the differentially expressed miRNAs and differentially expressed mRNAs were predicted using TargetScan (http://www.targetscan.org) [14], miRWalk (http://129.206.7.150/) and miRDB25378301. The interactions between the differentially expressed miRNAs and differentially expressed lncRNAs were predicted using the RNAhybrid program [15]. Then, the DEGs were merged with the target genes. The co-expressions of miRNAs, lncRNAs and mRNAs were selected for further analysis. The interactions of the lncRNAs-mRNAs-miRNAs were predicted using DIANA-TarBase v7.0 [16], and the threshold was a prediction score of >0.5. Cytoscape software (version 3.40) was used to visualize the regulatory network [17].

### Gene function analysis

Gene Ontology (GO) enrichment analysis of the differentially expressed genes was implemented with DAVID (http://david.abcc.ncifcrf.gov/). The GO terms included the three criteria: molecular function (MF), cellular component (CC), and biological process (BP). The P-values less than 0.05 were considered significantly enriched by the differentially expressed genes [18]. The Kyoto Encyclopaedia of Genes and the Genomes (KEGG) are database resources for understanding high-level functions and the effects of the biological system (http://www.genome.jp/kegg/)[19]. DAVID was also used to test the statistical enrichment of the genes or target genes of the miRNAs with differential expressions in the KEGG pathways [20]. The cutoff threshold was set as p-value<0.05.

### Protein-Protein Interaction (PPI) network construction and analysis

The protein-protein interaction network was constructed by using the STRING online database. The PPI pairs with selected larger scores were used to construct the PPI network[21. The transcription factors were annotated by using TF checkpoint (doi: 10.1093/bioinformatics/btt432). Then, the regulatory relationship between genes were visualized by using Cytoscape software (version 3.4.0) and analysed through topological property of computing network including the degree distribution of network by using CentiScaPe app [22]. And the gene with a degree>5 was defined as hub gene in the regulatory network according to the previous study [23].

### TCGA Datasets Analysis

TCGA is a platform for researchers to download and assess free public datasets (https://cancergenome.nih.gov/)[24]. In the present study, we verified the expression of hub genes in TCGA datasets to improve the reliability of our analysis.

## Results

### Identification of Differentially Expressed mRNAs, miRNAs and lncRNAs

The results show that 60 miRNAs, 1578 mRNAs and 128 lncRNAs were differentially expressed in prostate cancer (Tables S1,S2). The analysis of GSE64318 dataset: 60 miRNAs in PCa tumour tissues compared with normal prostate tissues were identified, including 35 up-regulated miRNAs and 25 down-regulated miRNAs. The analysis of GSE46602 dataset:1578 mRNAs including 583 up-regulated and 995 down-regulated mRNAs were identified; 128 lncRNAs including 49 up-regulated and 79 down-regulated lncRNAs were identified. Of which, we selected the top 10 significantly up-regulated and down-regulated mRNAs, miRNAs and lncRNAs in each category and plotted their expression of heat-maps and volcano plot ( Tables 1,2,3)(Figure1).

**Table 1.** Top 10 significantly up-regulated and down-regulated miRNAs in GSE64318 dataset.

| **miRNA_ID** | **Down/Up** | **logFC** | **adj.P.Val** | **P.Value** |
| --- | --- | --- | --- | --- |
| hsa-miR-671-5p | Down | -1.58 | 0.003129 | 6.91E-05 |
| hsa-miR-923 | Down | -1.5 | 0.01131 | 5.23E-04 |
| hsa-miR-483-5p | Down | -1.47 | 0.027441 | 2.04E-03 |
| hsa-miR-92b | Down | -1.42 | 0.004019 | 1.08E-04 |
| hsa-miR-150* | Down | -1.35 | 0.002095 | 2.81E-05 |
| hsa-miR-498 | Down | -1.31 | 0.007999 | 2.63E-04 |
| hsa-miR-933 | Down | -1.3 | 0.000927 | 6.46E-06 |
| hsa-miR-765 | Down | -1.3 | 0.001072 | 9.14E-06 |
| hsa-miR-563 | Down | -1.24 | 0.006355 | 1.94E-04 |
| hsa-miR-665 | Down | -1.21 | 0.003129 | 6.32E-05 |
| hsa-miR-221* | Up | 1.46 | 0.009099 | 3.21E-04 |
| hsa-miR-454 | Up | 1.46 | 0.029644 | 2.31E-03 |
| hsa-miR-26b | Up | 1.47 | 0.034161 | 3.16E-03 |
| hsa-let-7a | Up | 1.5 | 0.034161 | 3.02E-03 |
| hsa-miR-203 | Up | 1.51 | 0.003856 | 9.86E-05 |
| hsa-miR-148a | Up | 1.62 | 0.004387 | 1.23E-04 |
| hsa-miR-183 | Up | 1.63 | 0.009145 | 3.34E-04 |
| hsa-miR-200a | Up | 1.72 | 0.01323 | 7.09E-04 |
| hsa-miR-1 | Up | 1.92 | 0.034161 | 3.02E-03 |
| hsa-let-7f | Up | 2.01 | 0.024917 | 1.79E-03 |

**Table 2.** Top 10 significantly up-regulated and down-regulated mRNAs in GSE64318 dataset.

| **Gene Symbol** | **Down/Up** | **logFC** | **adj.P.Val** | **P.Value** |
| --- | --- | --- | --- | --- |
| CD177 | Down | -6.06431 | 2.03E-09 | 1.89E-12 |
| NEFH | Down | -5.84333 | 3.13E-05 | 3.34E-07 |
| SLC14A1 | Down | -5.45679 | 8.83E-10 | 7.15E-13 |
| KRT5 | Down | -5.10424 | 2.09E-10 | 1.03E-13 |
| KRT15 | Down | -4.99578 | 1.56E-10 | 7.00E-14 |
| TRIM29 | Down | -4.96486 | 2.68E-11 | 7.36E-15 |
| TP63 | Down | -4.90695 | 7.76E-11 | 2.98E-14 |
| JUP | Down | -4.73608 | 1.03E-09 | 9.03E-13 |
| NEFH | Down | -4.57684 | 5.07E-05 | 6.18E-07 |
| DST | Down | -4.55381 | 4.33E-08 | 7.60E-11 |
| CXCL14 | Up | 3.356898 | 2.96E-03 | 1.22E-04 |
| SIM2 | Up | 3.518509 | 2.60E-08 | 3.80E-11 |
| F5 | Up | 3.592734 | 1.63E-03 | 5.63E-05 |
| LUZP2 | Up | 3.789194 | 8.41E-05 | 1.21E-06 |
| PCA3 | Up | 3.822882 | 1.28E-04 | 2.08E-06 |
| AMACR | Up | 3.830373 | 7.49E-07 | 2.88E-09 |
| GLYATL1 | Up | 3.928912 | 8.57E-07 | 3.45E-09 |
| AMACR | Up | 4.307398 | 3.25E-07 | 1.02E-09 |
| DLX1 | Up | 4.410439 | 5.09E-06 | 3.35E-08 |
| PCA3 | Up | 4.558841 | 7.89E-06 | 5.87E-08 |

**Table 3.** Top 10 significantly up-regulated and down-regulated lncRNAs in GSE46602.

| **Gene Symbol** | **Location** | **adj.P.Val** | **P.Value** | **logFC** | **Down/Up** |
| --- | --- | --- | --- | --- | --- |
| LINC00844 | chr10:58999518-59001617 | 5.83E-09 | 6.18E-12 | -4.45143 | Dwon |
| MIR205HG | chr1:209428820-209432552 | 4.37E-13 | 5.59E-17 | -3.34821 | Dwon |
| RP11-44F21.5 | chr4:75081702-75084717 | 2.14E-07 | 6.06E-10 | -3.01543 | Dwon |
| RP11-680G24.5 | chr16:15018106-15020488 | 1.49E-05 | 1.28E-07 | -2.57293 | Dwon |
| RP11-203B7.2 | chr4:146052604-146056762 | 2.96E-07 | 9.10E-10 | -2.25844 | Dwon |
| MAGI2-AS3 | chr7:79452957-79471208 | 1.29E-07 | 3.08E-10 | -1.94617 | Dwon |
| LSAMP-AS1 | chr3:116360024-116370085 | 4.05E-06 | 2.48E-08 | -1.94409 | Dwon |
| LINC00937 | chr12:8356963-8390752 | 1.63E-04 | 2.83E-06 | -1.93046 | Dwon |
| LINC00883 | chr3:107240692-107326964 | 3.22E-04 | 6.79E-06 | -1.86947 | Dwon |
| PGM5-AS1 | chr9:68355164-68357866 | 4.00E-02 | 4.20E-03 | -1.86179 | Dwon |
| RP5-1092A3.4 | chr1:28239509-28241453 | 1.16E-01 | 2.21E-02 | 1.132616 | Up |
| LINC01023 | chr5:108727825-108728261 | 9.06E-04 | 2.61E-05 | 1.13526 | Up |
| LINC01128 | chr1:827591-859446 | 4.49E-02 | 5.00E-03 | 1.199525 | Up |
| BAIAP2-AS1 | chr17:81029133-81034719 | 1.48E-03 | 4.96E-05 | 1.201015 | Up |
| RP11-48B3.4 | chr8:80541300-80543104 | 1.15E-02 | 7.49E-04 | 1.284279 | Up |
| RP11-339B21.15 | chr9:128305161-128305515 | 6.32E-05 | 8.40E-07 | 1.310892 | Up |
| GABPB1-AS1 | chr15:50354174-50358312 | 8.11E-04 | 2.26E-05 | 1.367447 | Up |
| RP11-391M1.4 | chr3:40466466-40468587 | 7.89E-06 | 5.87E-08 | 1.489649 | Up |
| RP11-295G20.2 | chr1:231522517-231528556 | 7.32E-04 | 1.96E-05 | 1.673082 | Up |
| LINC00992 | chr5:117415512-117579744 | 1.90E-04 | 3.44E-06 | 2.288701 | Up |

**Figure 1.**
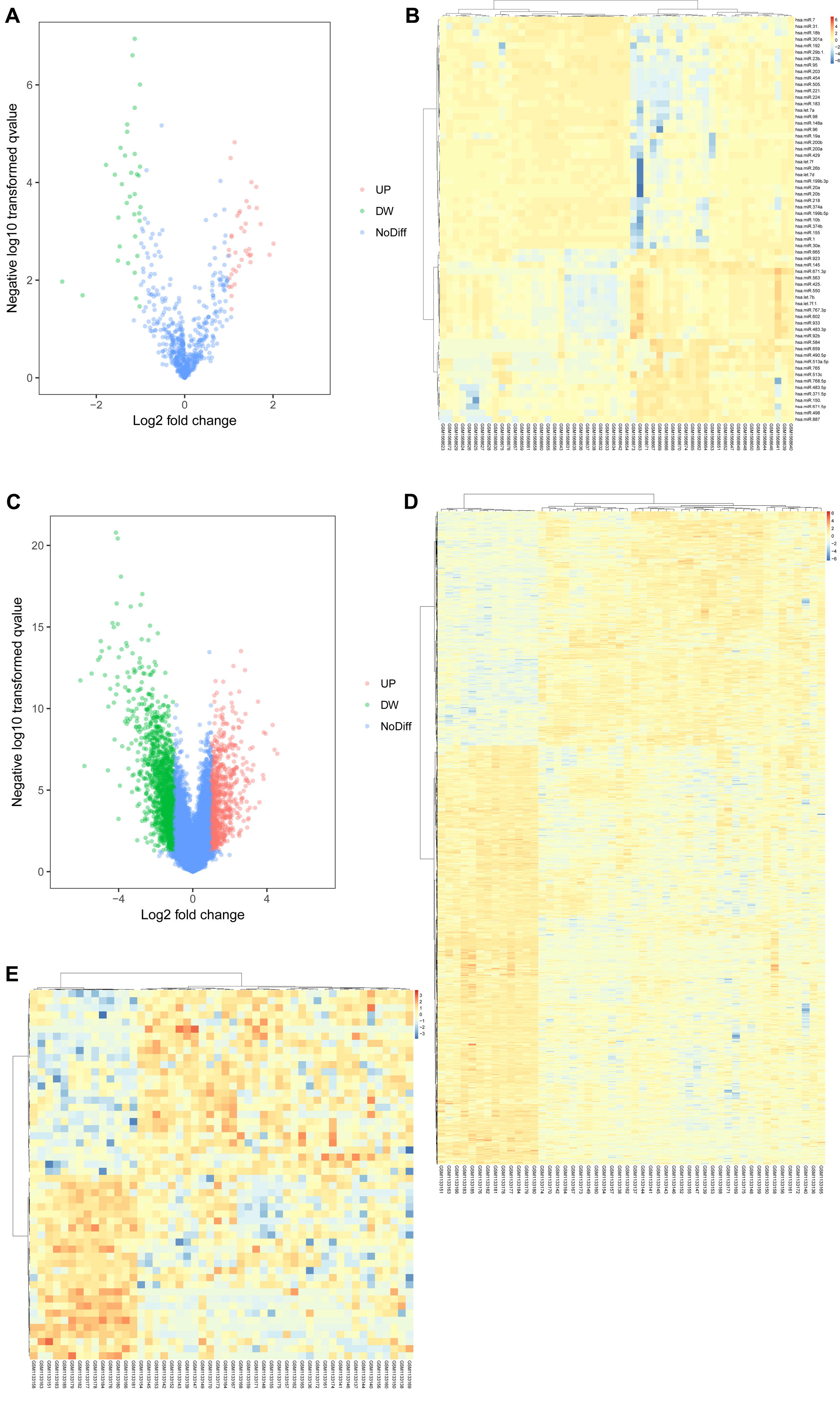
The differentially expressed genes in prostate cancer (PCa) and normal prostate tissues. (A. Volcano plot of the differentially expressed miRNAs. B. heatmap of the differentially expressed miRNAs. C. Volcano plot of the differentially expressed mRNAs. D. Heatmap of the differentially expressed mRNAs. E. Heatmap of differentially expressed lncRNAs)

### mRNAs-miRNAs-lncRNAs network analysis

miRNAs–mRNAs regulatory network was constructed. Interaction analysis show that 33 differentially expressed miRNAs targeted 143 mRNAs up or down (Figure 2). In detail, 25 over-expressed miRNAs down-regulate 104 mRNAs. 8 down-expressed miRNAs up-regulate 15 mRNAs. Subsequently, we reconstruct the lncRNAs ‑miRNAs–mRNAs network after selecting the pre-treated data by P-values <0.05 and fold change>2 (Figure 3). In the network, there are 5 miRNA nodes, 13 lncRNA nodes, 45 mRNA nodes. In detail, 9 down-expressed lncRNAs up-regulate 19 miRNAs, 5 over-expressed miRNAs down-regulate 44 mRNAs.

**Figure 2.**
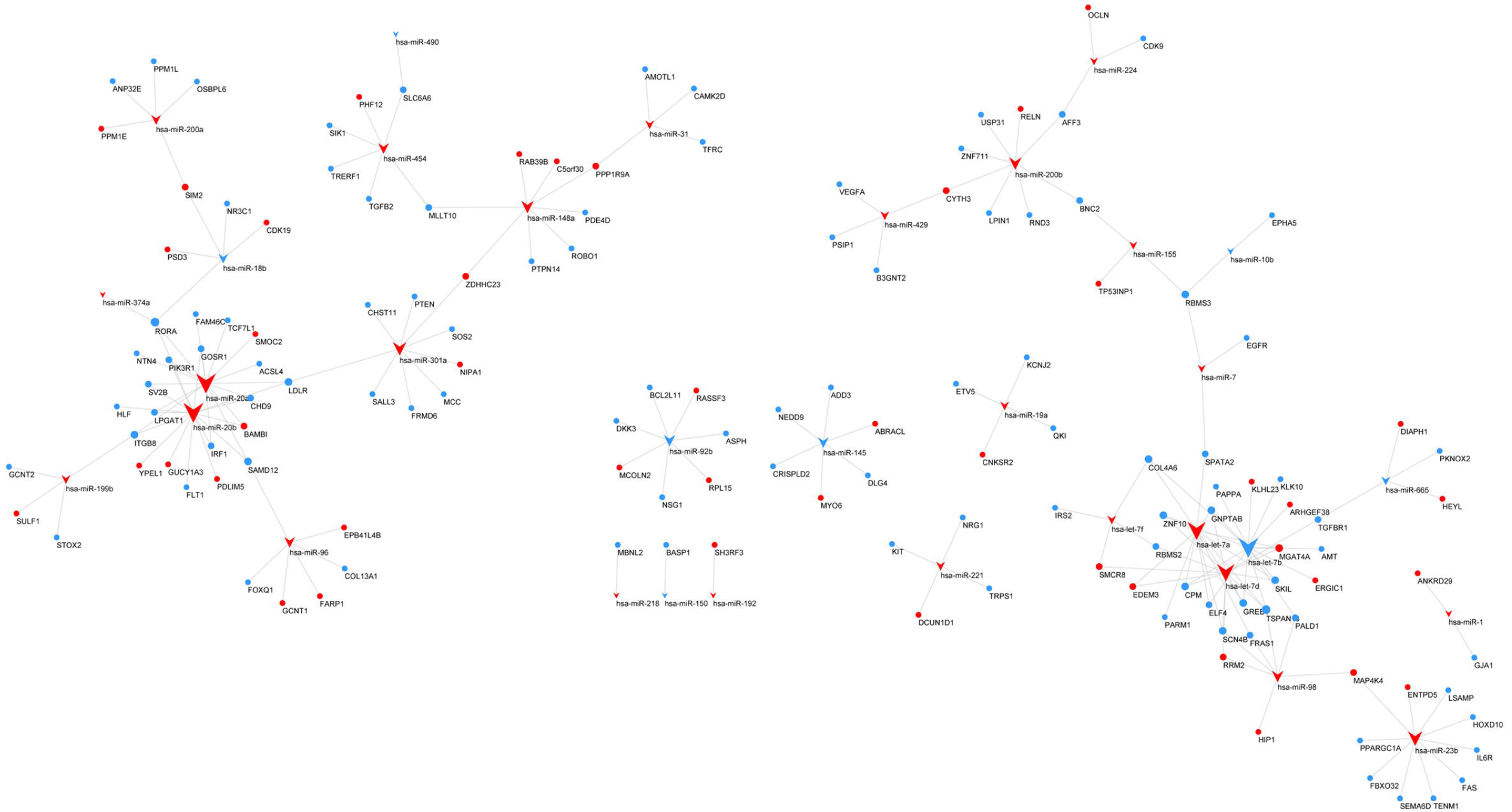
The miRNAs–mRNAs regulatory network. miRNAs are indicated with arrows, and mRNAs are indicated with circles. The colour red represents high expression, and green represents low expression.

**Figure 3.**
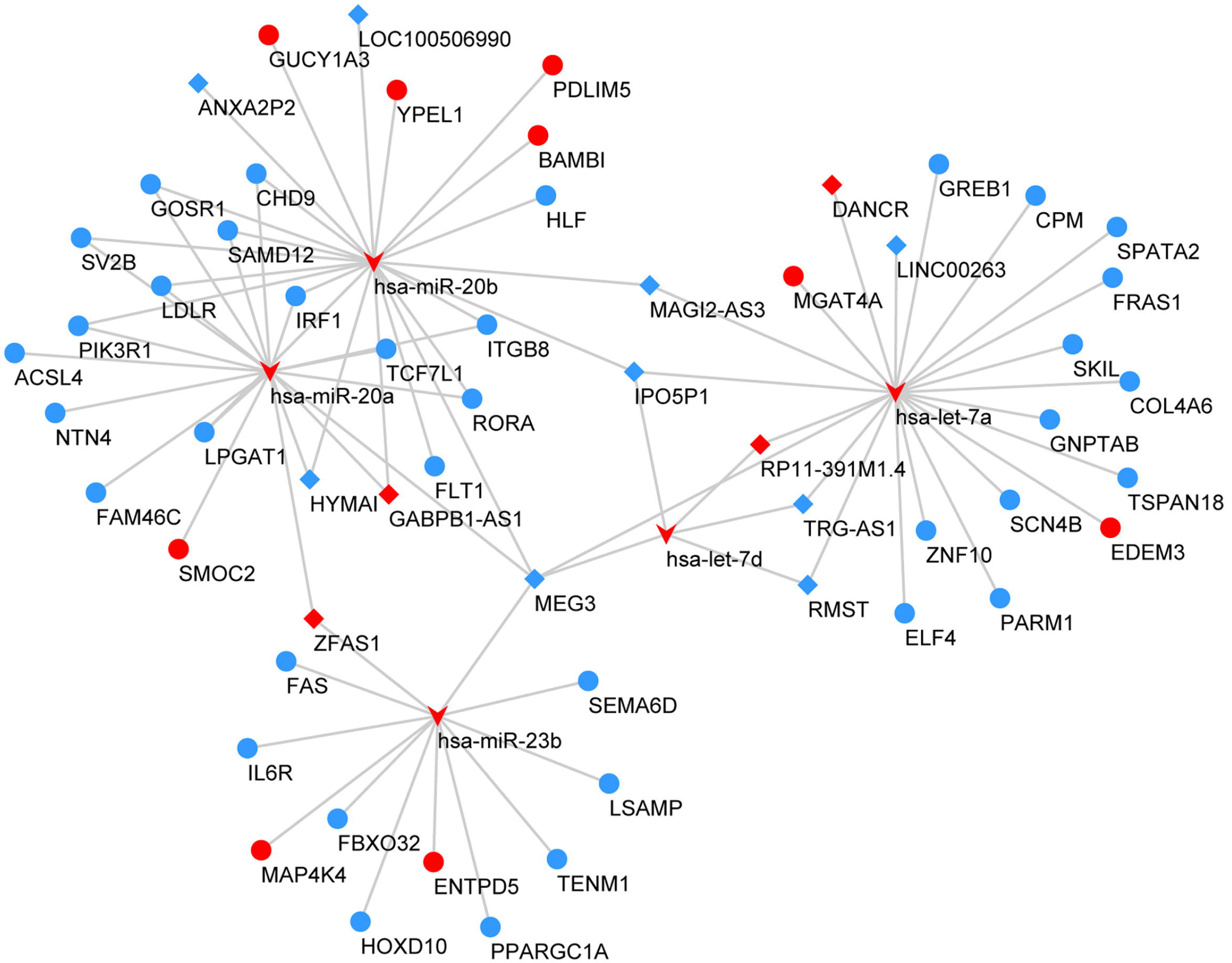
The regulatory network of the mRNAs-miRNAs-lncRNAs. mRNAs, miRNAs and lncRNAs are indicated by the circles, the arrow and diamond shape, respectively. The colour red represents high expression, and green represents low expression.

### Function enrichment analysis of DEGs

GO enrichment analyses for DEGs based on mRNAs-miRNAs-lncRNAs network were performed. The top 10 most significant GO terms of each group were shown. On the MF level, the DEGs were mainly enriched in sequence-specific DNA binding. On the CC level, the DEGs were mainly enriched in nucleus, cytoplasm and plasma membrane. On the BP level, the DEGs were mainly enriched in positive regulation of transcription from RNA polymerase II promoter, cell proliferation and cell migration. The top 20 most significant KEGG pathway terms are shown. The genes were mainly enriched in PI3K-Akt signalling pathway, pathways in cancer and focal adhesion signalling pathway (Figure 4) (Tables S3).

**Figure 4.**
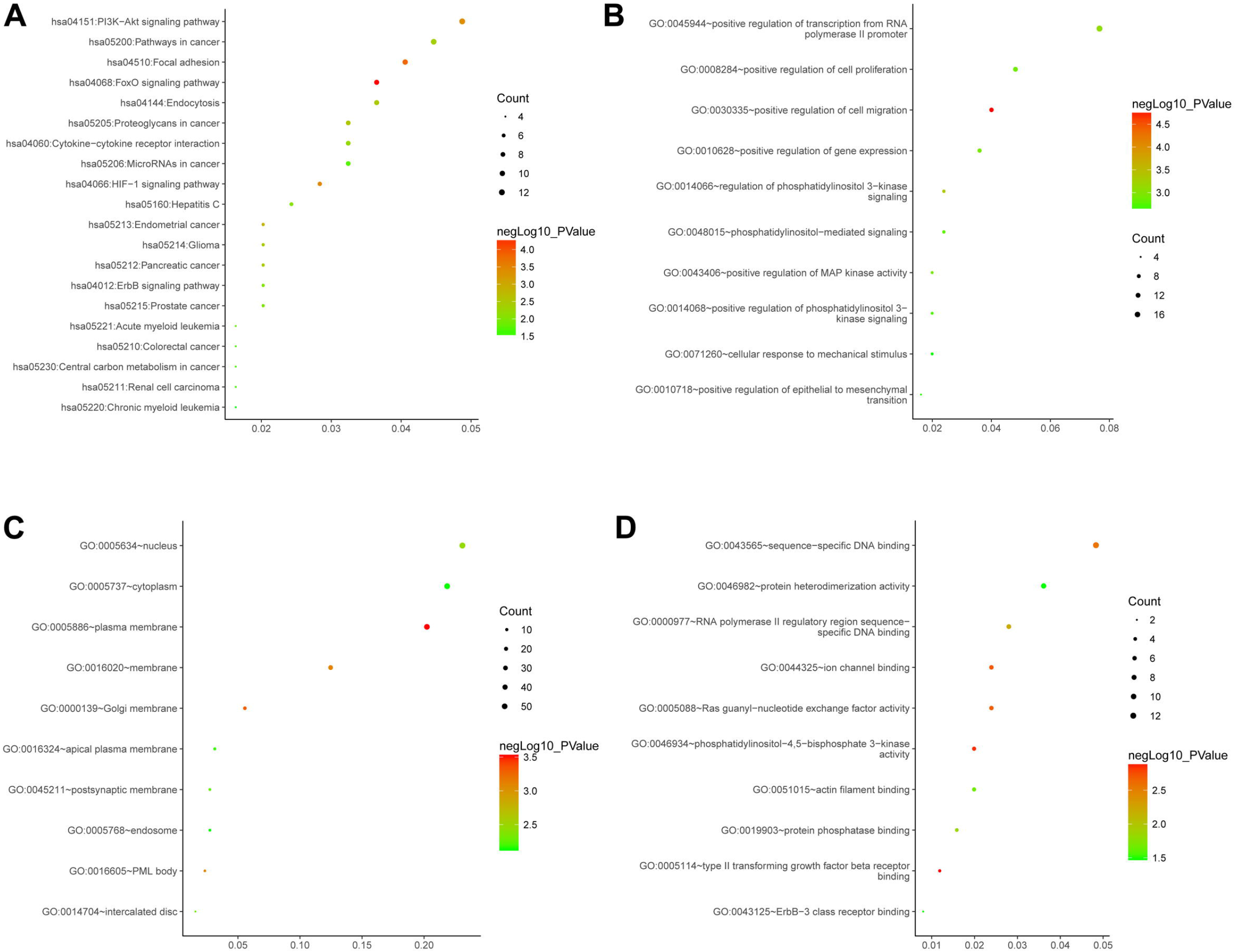
Gene ontology (GO) and Kyoto Encyclopedia of Genes and Genome (KEGG) enrichment analysis. (A. The top 20 most significant KEGG pathway terms. B. The top 10 most significant changes in the GO biological process. C. The top 10 most significant changes in the GO cellular component. D. The top 10 most significant changes in the GO molecular function.)

### Protein-protein interaction network (PPI) analysis

Using the STRING online database and the Cytoscape software, a total of 128 DEGs of the 1578 DEGs were mapped into the PPI network complex. In this network, 32 nodes were chosen as hub nodes and included 2 TFs, 14 miRNAs, and 16 mRNAs; the results are presented in Table 4 and Figure 5. Moreover, we found that 6 mRNAs (i.e., EGFR, VEGFA, PIK3R1, DLG4, TGFBR1 and KIT) exhibited higher degrees (>10) and miRNA-mRNA pairs. These findings suggest that these nodes may play important roles in the development of PCa.

**Table 4.** The topology parameter of the hub genes (degree>5) in the miRNA-mRNA-TF regulatory network.

| **Gene** | **Closeness** | **Betweenness** | **Degree** |
| --- | --- | --- | --- |
| EGFR | 0.00212766 | 4046.701286 | 22 |
| VEGFA | 0.002159827 | 4331.82023 | 20 |
| hsa-miR-20b | 0.001855288 | 2752.2272 | 16 |
| hsa-miR-20a | 0.001821494 | 2973.867274 | 16 |
| hsa-let-7b | 0.001481481 | 2436.558748 | 16 |
| hsa-let-7a | 0.001582278 | 1383.644882 | 14 |
| hsa-let-7d | 0.00174216 | 2553.049636 | 13 |
| PIK3R1 | 0.002057613 | 2045.442867 | 12 |
| DLG4 | 0.001869159 | 3167.065846 | 12 |
| TGFBR1 | 0.001976285 | 3691.770395 | 11 |
| KIT | 0.001904762 | 1517.550434 | 11 |
| hsa-miR-23b | 0.001587302 | 1375.820161 | 10 |
| PTEN | 0.001923077 | 1345.754047 | 10 |
| hsa-miR-301a | 0.001610306 | 1995.023424 | 9 |
| LDLR | 0.001960784 | 1888.001784 | 9 |
| FLT1 | 0.001930502 | 1179.255452 | 9 |
| hsa-miR-200b | 0.001445087 | 1353.57487 | 8 |
| hsa-miR-148a | 0.00140647 | 1235.527073 | 8 |
| PPARGC1A | 0.001801802 | 1571.190602 | 8 |
| BCL2L11 | 0.001834862 | 1606.829419 | 8 |
| NR3C1* | 0.00177305 | 733.3472529 | 7 |
| NRG1* | 0.001872659 | 1359.250724 | 7 |
| hsa-miR-98 | 0.001618123 | 802.5761652 | 7 |
| hsa-miR-92b | 0.001506024 | 1655.33632 | 7 |
| GJA1 | 0.001745201 | 1149.048086 | 7 |
| hsa-miR-96 | 0.001335113 | 1416.966025 | 6 |
| hsa-miR-454 | 0.001420455 | 1854.243032 | 6 |
| hsa-miR-145 | 0.001461988 | 1257.608267 | 6 |
| TFRC | 0.001821494 | 445.8511305 | 6 |
| ITGB8 | 0.001808318 | 2026.300029 | 6 |
| EPHA5 | 0.001689189 | 845.4402231 | 6 |
| CAMK2D | 0.001620746 | 1045.62274 | 6 |

\* transcription factors (TFs)

**Figure 5.**
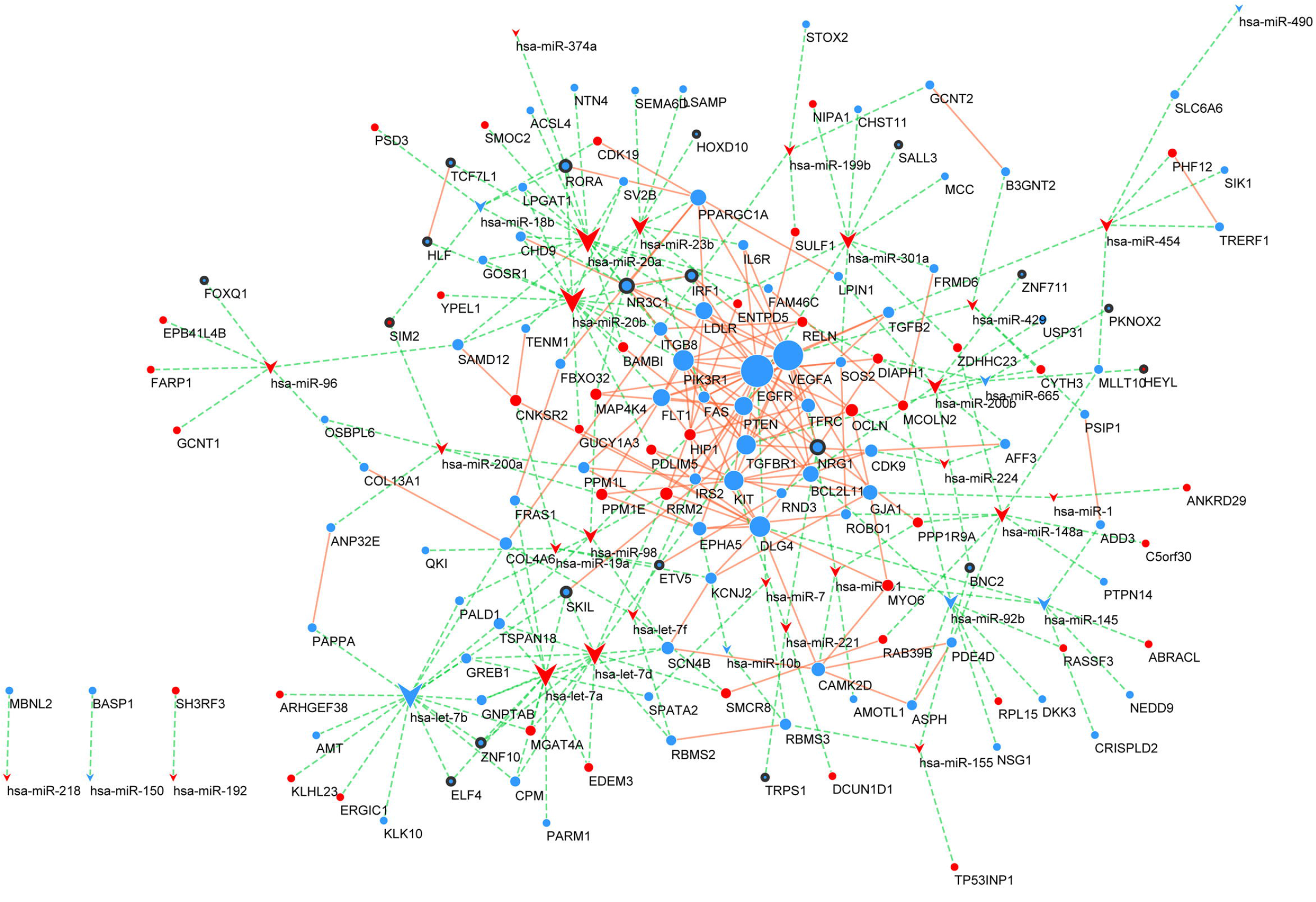
Protein-protein interaction (PPI) network. mRNAs and miRNAs are indicated with circle sand arrows. TFs are indicated with black-traced circles. The green dotted line indicates the interaction of the miRNAs-mRNAs. The orange line indicates the relationship of the interaction of the proteins. The colour red represents high expression, and green represents low expression.

### TCGA Datasets Analysis

The TCGA dataset analyses were performed to demonstrate that the aberrant expression of the hub genes, including EGFR, VEGFA, PIK3R1, DLG4, TGFBR1 and KIT, were significantly different between PCa and normal prostate tissues (Figure 6).

**Figure 6.**
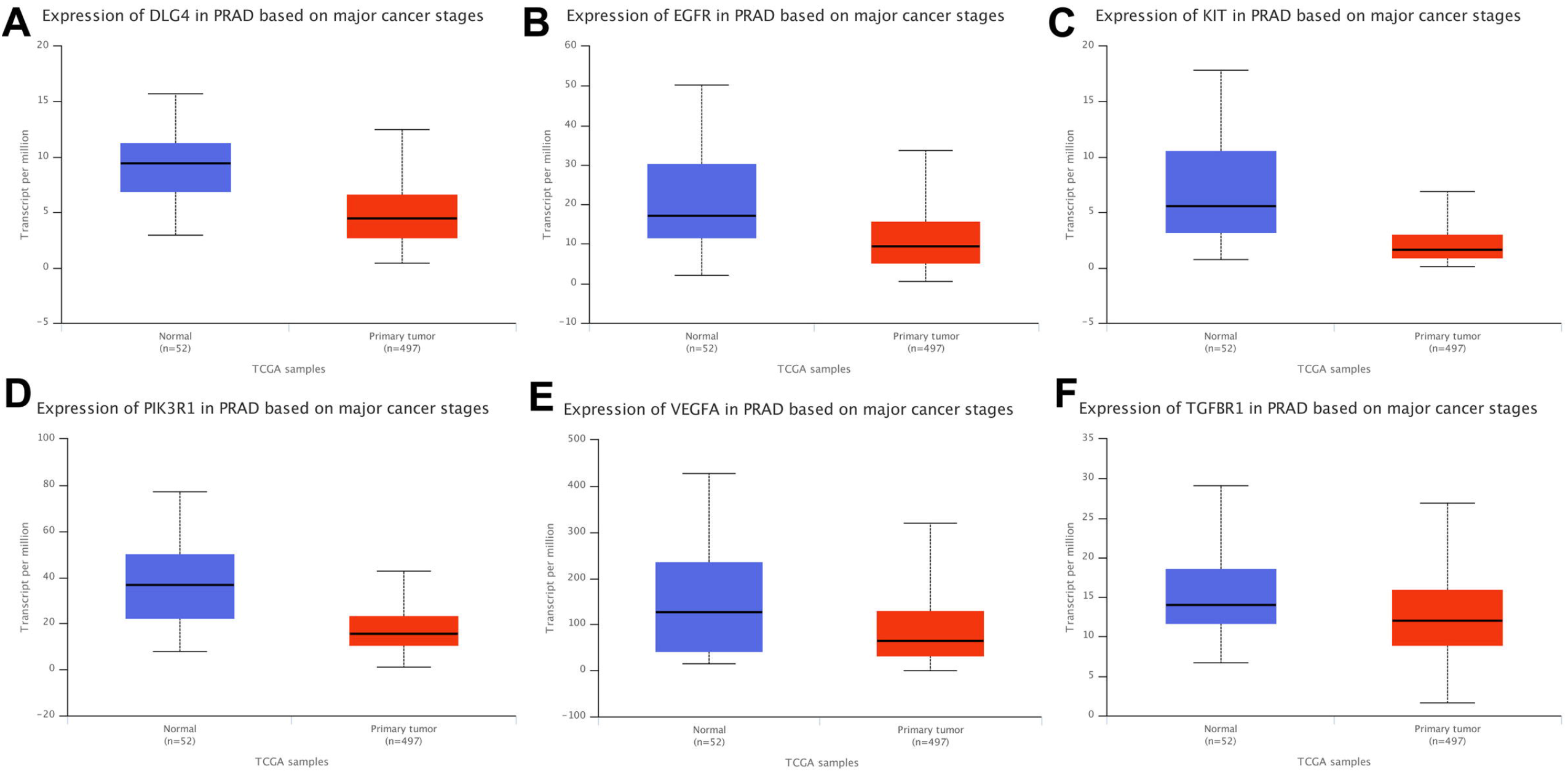
TCGA dataset analysis. (A. The expression of DLG4 in PCa. B. The expression of EGFR in PCa. C. The expression of KIT in PCa. D. The expression of PIK3R1 in PCa. E. The expression of VEGFA in PCa. F. The expression of TGFBR1
in PCa)

## Discussion

Prostate cancer (PCa) has become a public health issue of great concern worldwide [25]. However, the molecular mechanisms involved in the progress of PCa are still unclear. Therefore, it is very crucial to study the mechanism and to identify molecular targets for diagnosis and treatment. In this study, we performed a comprehensive bioinformatics analysis and retrieved the mRNAs, miRNAs, lncRNAs and TFs in the interaction network and revealed the key genes relevant to prostate cancer.

MiRNA expressions in tumour tissues often differ from that of normal tissue, and the differential expression affects the occurrence, development and prognosis of tumours [26, 27]. Studies have confirmed that miRNA expression profiles can be used as biomarkers for the early detection, classification and prognosis of tumours [28, 29]. lncRNAs act as miRNAs sponges and can regulate miRNAs abundance and compete with mRNAs for the binding of miRNAs [30]. By constructing a mRNA-miRNA-lncRNA network, we found that the aberrant expression of lncRNAs led to the abnormal expression of 5 miRNAs (i.e., has-miR-20a, has-miR-20b, has-miR-23b, has-let-7a and has-let-7d) in PCa and thus regulated the expression of the target mRNAs. A previous study demonstrated that 5 miRNAs regulate the development of prostate cancer by targeting specific mRNAs and play an important role in the regulation of prostate cancer [31,32,33]. The involvements of key lncRNAs including HYMAI, MEG3, IPO5P1, MAG12-AS3, RMST and TRG-AS1 are important. Among them, MEG3 is an important tumour suppressor gene that inhibits cell proliferation and induces apoptosis in PCa [34]. The finding of these lncRNAs suggests potential diagnostic and therapeutic targets for PCa.

Next, we conducted a functional enrichment analysis of the DEGs based on the mRNAs-miRNAs-lncRNAs network. We found that the DEGs were mainly enriched in the nucleus and cytoplasm, were involved in the regulation of transcription, were related to the sequence-specific DNA binding, and participated in the regulation of the PI3K-Akt signalling pathway; this pathway is related to cancer and the focal adhesion signalling pathway. To further analysed the key genes related to PCa, we constructed a PPI network. More significantly, we found that the transcription factor NR3C1 was a hub gene that participated in the regulation of the expression of multiple miRNAs (has-miR-20a, has-miR-20b, has-miR-23b). Puhr M et al assessed NR3C1 expression and the functional significance in tissues from PCa and found that it is a key factor for the development of PCa [35]. Moreover, we found that its regulatory function is similar to MEG3 and its down-regulation leads to increased expression of miRNAs (has-miR-20a, has-miR-20b, and has-miR-23b). Based on these results, we speculate that there may be some regulatory relationship between NR3C1 and MEG3.

Finally, through PPI and TCGA dataset analyses, we found that 6 mRNAs (i.e., EGFR, VEGFA, PIK3R1, DLG4, TGFBR1 and KIT) had higher degrees and miRNA-mRNA pairs. These expressions were lower and were significantly different between PCa and normal prostate tissues. EGFR belongs to a family of cell membrane receptor tyrosine kinases and is a key factor in tumour cell growth and invasion [36, 37]. Previous studies have demonstrated that abnormal expression of EGFR and its downstream signalling contribute to disease progression in PCa [38]. The downregulation of the EGFR is associated with enhanced signalling [39], which can lead to the development of cancer [40]. VEGFA is a mitogen with high endothelial cell specificity and plays a major regulatory role in the development of PCa [41]. A previous study found that PIK3R1, TGFBR1 and KIT might have clinical utilities in distinguishing PCa [42,43,44]. We also found that 6 mRNAs (EGFR, VEGFA, PIK3R1, DLG4, TGFBR1 and KIT) were associated with the abnormal expression of 5 miRNAs (has-miR-20a, has-miR-20b, has-miR-23b, has-let-7a and has-let-7d) in PCa. In conclusion, the hub genes that we identified might play crucial roles in PCa.

## Conclusion

We constructed and analysed mRNAs, miRNAs, lncRNAs, and TF interaction networks to reveal the key genes in prostate cancer. We found that 5 miRNAs (has-miR-20a, has-miR-20b, has-miR-23b, has-let-7a and has-let-7d), 6 lncRNAs (HYMAI, MEG3, IPO5P1, MAG12-AS3, RMST and TRG-AS1),6 mRNAs (EGFR, VEGFA, PIK3R1, DLG4, TGFBR1 and KIT) and 2TFs (NR3C1,NRG1) play important regulatory roles in the interaction network. The expression levels of EGFR, VEGFA, PIK3R1, DLG4, TGFBR1 and KIT were \significantly different between PCa and normal tissues. Further research is needed to specify the molecular mechanism of these hub genes in PCa.

## Acknowledgements

The authors would like to thank the study subjects and research personnel for their involvement in the study. The authors would also like to thank Professor Xiaoke Hao and Professor Yueyun Ma for their excellent theoretical assistance.

## Author contributions

Yun Ye designed the overall project and wrote the manuscript. Yun Ye and Su-Liang Li collected and analysed the data. Sheng-Yu Wang provided the analytical software.

## REFERENCES

[1]. Ferlay J, Shin HR, Bray F, et al. Estimates of worldwide burden of cancer in 2008:GLOBOCAN 2008. Int J Cancer.2010;127(12): 2893–2917.

[2]. Siegel RL, Miller KD, Jemal A. Cancer Statistics, 2017. CA Cancer J Clin. 2017 ; 67(1):7–30.

[3]. Larson SR, Zhang X, Dumpit R, et al. Characterization of osteoblastic and osteolytic proteins in prostate cancer bone metastases. Prostate. 2013; 73(9): 932–940.

[4]. Rajpar S, Fizazi K. Bone targeted therapies in metastatic castration-resistant prostate cancer. Cancer J. 2013;19(1):66–70.

[5]. Guttman M, Rinn JL. Modular regulatory principles of large non-coding RNAs. Nature. 2012;482(7385):339–346.

[6]. Chen L, Yao H, Wang K, Liu X. Long Non-Coding RNA MALAT1 Regulates ZEB1 Expression by Sponging miR-143-3p and Promotes Hepatocellular Carcinoma Progression. J Cell Biochem. 2017;118(12):4836–4843.

[7]. Ali Sobhi Afshar, Joseph Xu, John Goutsias. Integrative Identification of Deregulated MiRNA/TF-Mediated Gene Regulatory Loops and Networks in Prostate Cancer. PLoS One. 2014; 9(6): e100806.

[8]. Xu H, He JH, Xiao ZD, et al. Liver-enriched transcription factors regulate microRNA-122 that targets CUTL1 during liver development. Hepatology. 2010;52(4):1431–1442.

[9]. Vogelstein B, Papadopoulos N, Velculescu VE, et al. Cancer genome landscapes. Science. 2013; 339(6127): p. 1546–1558.

[10]. Guo Y, Bao Y, Ma M, Yang W. Identification of Key Candidate Genes and Pathways in Colorectal Cancer by Integrated Bioinformatical Analysis. Int J Mol Sci. 2017;18(4). pii: E722.

[11]. Edgar R, Domrachev M, Lash AE. Gene Expression Omnibus: NCBI gene expression and hybridization array data repository. Nucleic Acids Res. 2002; 30(1): 207–210.

[12]. Irizarry RA, Bolstad BM, Collin F, et al. Summaries of Affymetrix GeneChip probe level data. Nucleic Acids Res. 2003;31(4):e15.

[13]. Diboun I, Wernisch L, Orengo CA, Koltzenburg M. Microarray analysis after RNA amplification can detect pronounced differences in gene expression using limma. BMC Genomics. 2006;7:252.

[14]. Lewis BP, Burge CB, Bartel DP. Conserved seed pairing, often flanked by adenosines, indicates that thousands of human genes are microRNA targets. Cell.2005; 120 (1): 15–20.

[15]. Rehmsmeier M, Steffen P, Hochsmann M, Giegerich R. Fast and effective prediction of microRNA/target duplexes. RNA. 2004;10(10):1507–1517.

[16]. Vlachos IS, Paraskevopoulou MD, Karagkouni D, et al. DIANA-TarBase v7.0: indexing more than half a million experimentally supported miRNA:mRNA interactions. Nucleic Acids Res. 2015;43(Database issue):D153–159.

[17]. Bindea G, Mlecnik B, Hackl H, Charoentong P, et al. ClueGO: a Cytoscape plug-in to decipher functionally grouped gene ontology and pathway annotation networks. Bioinformatics.2009; 25(8): 1091–1093.

[18]. Ashburner M, Ball CA, Blake JA, et al. Gene ontology: tool for the unification of biology. The Gene Ontology Consortium. Nat Genet. 2000;25(1): 25–29.

[19]. Kanehisa M, Goto S. KEGG: kyoto encyclopedia of genes and genomes. Nucleic Acids Res. 2000; 28(1): 27–30.

[20]. Huang da W, Sherman BT, Lempicki RA. Lempicki, Systematic and integrative analysis of large gene lists using DAVID bioinformatics resources. Nat Protoc. 2009;4(1): 44–57.

[21]. Szklarczyk D, Franceschini A, Wyder S, et al. STRING v10: protein-protein interaction networks, integrated over the tree of life. Nucleic Acids Res. 2015; 43(Database issue): D447–452.

[22]. Scardoni G, Tosadori G, Faizan M, et al. Biological network analysis with CentiScaPe: centralities and experimental dataset integration. Version 2. F1000Res. 2014;3:139.

[23]. Han JD, Bertin N, Hao T, et al. Evidence for dynamically organized modularity in the yeast protein-protein interaction network. Nature. 2004;430(6995):88–93.

[24]. Tomczak K, Czerwinska P, Wiznerowicz M. The Cancer Genome Atlas (TCGA): an immeasurable source of knowledge. Contemp Oncol (Pozn). 2015;19(1A): A68–77.

[25]. Roy Mano, James Eastham, Ofer Yossepowitch. The Very High Risk Prostate Cancer – a Contemporary Update. Prostate Cancer Prostatic Dis. 2016; 19(4): 340–348.

[26]. Song C, Chen H, Wang T, et al. Expression profile analysis of micro RNAs in prostate cancer by next-generation sequencing. Prostate. 2015; 75(5): 500–516.

[27]. Avril S. [micro RNA expression in breast development and breast cancer]. Pathologe. 2013;34 Suppl 2: 195–200.

[28]. Ferracin M, Veronese A, Negrini M. Micromarkers: miRNAs in cancer diagnosis and prognosis. Expert Rev Mol Diagn. 2010;10(3):297–308.

[29]. Søkilde R, Vincent M, Møller AK, et al. Efficient identification of miRNAs for classification of tumor origin. J Mol Diagn.2014;16(1):106–115.

[30]. Wang JB, Liu FH, Chen JH, et al. Identifying survival-associated modules fromthe dysregulated triplet network in glioblastoma multiforme. J Cancer Res Clin Oncol. 2017;143(4):661–671.

[31]. Liu DF, Wu JT, Wang JM, et al. MicroRNA expression profile analysis reveals diagnostic biomarker for human prostate cancer. Asian Pac J Cancer Prev. 2012;13(7):3313–3317.

[32]. Cai S, Chen R, Li X,et al. Downregulation of microRNA-23a suppresses prostate cancer metastasis by targeting the PAK6-LIMK1 signaling pathway. Oncotarget. 2015;6(6):3904–3917.

[33]. Wagner S, Ngezahayo A, Murua Escobar H, Nolte I. Role of miRNA let-7 and its major targets in prostate cancer. Biomed Res Int. 2014;2014:376326.

[34]. Luo G, Wang M, Wu X, et al. Long Non-Coding RNA MEG3 Inhibits Cell Proliferation and Induces Apoptosis in Prostate Cancer. Cell Physiol Biochem. 2015;37(6):2209–2220.

[35]. Puhr M, Hoefer J, Eigentler A, et al. The glucocorticoid receptor is a key player for prostate cancer cell survival and a target for improved antiandrogen therapy. Clin Cancer Res. 2018;24(4):927–938.

[36]. Lim SO, Li CW, Xia W, et al. EGFR Signaling Enhances Aerobic Glycolysis in Triple-Negative Breast Cancer Cells to PromoteTumor Growth and Immune Escape. Cancer Res. 2016;76(5):1284–1296.

[37]. Daizumoto K, Yoshimaru T, Matsushita Y, et al. A DDX31/mutant-p53/EGFR axis promotes multistep progression of muscle invasive bladder cancer. Cancer Res. 2018 ; pii: canres.2528.2017.

[38]. Di Lorenzo G, Tortora G, D’Armiento FP, et al. Expression of epidermal growth factor receptor correlates with disease relapse and progression to androgen-independence in human prostate cancer. Clin Cancer Res. 2002; 8(11): 3438–3444.

[39]. Sorkin A, von Zastrow M. Endocytosis and signalling: intertwining molecular networks. Nat Rev Mol Cell Biol. 2009;10(9):609–622.

[40]. Roepstorff K, Grøvdal L, Grandal M, et al. Endocytic downregulation of ErbB receptors: mechanisms and relevance in cancer. Histochem Cell Biol. 2008; 129(5): 563–578.

[41]. Lee C, Whang YM, Campbell P, et al. Dual targeting c-met and VEGFR2 in osteoblasts suppresses growth and osteolysis of prostate cancer bone metastasis. Cancer Lett. 2018;414:205–213.

[42]. Hellwinkel OJ, Rogmann JP, Asong LE, et al. A comprehensive analysis of transcript signatures of the phosphatidylinositol-3 kinase/protein kinase B signal-transduction pathway in prostate cancer. BJU Int. 2008; 101(11):1454–1460.

[43]. Wen J, Li R, Wen X, et al. Dysregulation of cell cycle related genes and microRNAs distinguish the low-from high-risk of prostate cancer. Diagn Pathol. 2014;9:156.

[44]. Fonseca-Alves CE, Kobayashi PE, Palmieri C, Laufer-Amorim R. Investigation of c-KIT and Ki67 expression in normal, preneoplastic and neoplastic canine prostate. BMC Vet Res. 2017;13(1):380.

